# Neural stem cell quiescence is actively maintained by the epigenome

**DOI:** 10.1101/2025.05.14.653490

**Authors:** Anna Malkowska, Jan Ander, Andrea H. Brand

## Abstract

Homeostasis and repair of the nervous system is maintained by a population of resident neural stem cells (NSCs) retained in a state of reversible cell cycle arrest called quiescence. quiescent NSCs can resume proliferation in response to different physiological stimuli, such as diet or injury. Reactivation of NSCs requires changes in gene expression, much of which is regulated at the epigenomic level. We mapped, comprehensively and *in vivo*, the dynamic epigenomic changes in NSC chromatin during stem cell quiescence and reactivation in *Drosophila*. Contrary to expectations, we found that chromatin accessibility is increased in quiescent NSCs, remodelling extensively within both euchromatin and heterochromatin. Surprisingly, genes crucial for cell cycle progression are repressed whilst remaining within permissive H3K36me3-bound euchromatin. At the same time, genes necessary for cell-cell communication are derepressed by eviction of histone H1 and transition to a SWI/SNF-enriched active state. Our results reveal global expansion of accessible chromatin in quiescent NSCs without concomitant activation of transcription. Strikingly, this process reverses upon reactivation, indicating that opening of chromatin is a quiescence-specific event.

## Introduction

Many adult tissues retain a population of quiescent stem cells to mediate repair and maintain tissue homeostasis in response to the changing physiology. Quiescence is an actively maintained state of cell cycle arrest associated with differential transcription, epigenomic state and metabolism^1^. In the adult mammalian nervous system, quiescent neural stem cells (NSCs) are found in the subventricular zone of the lateral ventricles, the dentate gyrus of the hippocampus and the median eminence of the hypothalamus, where they contribute to neurogenesis in response to cues such as changing diet, exercise, pregnancy or injury^2–5^.

Quiescence is widely thought to be associated with chromatin condensation and therefore transcriptional and metabolic inactivity^6,7^. However, the quiescent state needs to be actively maintained, as quiescent cells have to retain their self-renewal and differentiation capabilities through cell cycle exit and resume proliferation in response to triggers whilst in communication with their niche^7,8^. A careful examination of quiescent stem cell models reveals a variety of chromatin changes dependent on the tissue of origin^9^. In muscle stem cells, quiescence is characterised by increased H4K20me3-mediated repression^10^. Quiescent lymphocytes and hair follicle stem cells, on the other hand, have lower levels of both active and repressive histone methylation^11–13^. Consequently, quiescent chromatin signature is tissue-specific and dependent on developmental context^9^ and, therefore, studying the chromatin of quiescent cells in their native environment is crucial. The sparseness of neural stem cells (NSCs) in the mammalian brain and the lack of well-defined genetic markers leads to a difficulty in profiling the genome of mammalian NSCs whilst still in their niche. As a result, studies investigating chromatin changes in quiescent and active NSCs have focused on chromatin accessibility^14,15^ instead of the full repertoire of chromatin proteins and histone modifications. As such, little is known about the specific epigenetic modifiers that might regulate mammalian NSC quiescence and reactivation *in vivo*.

In contrast to the mammalian nervous system, NSCs in *Drosophila melanogaster* are easily identifiable *in vivo* by well-defined genetic markers and the stereotypic developmental timeline of quiescence induction and reactivation. *Drosophila* NSCs share many features with their mammalian counterparts, including similar morphology, reliance on oxidative phosphorylation during proliferation, homologous quiescence factors such as Cyclin dependent kinase inhibitors, and Akt-dependent reactivation triggered by a dietary signal^16–20^. *Drosophila* neural stem cells in the embryonic and larval brain serve as a well-studied and more easily accessible natural quiescence model^21,22^. However, the understanding of how quiescent and active NSC gene signatures are established and maintained at the chromatin level and whether epigenomic pathways instruct quiescence induction and reactivation remains lacking.

The fundamental unit of chromatin is the nucleosome, which consists of a covalently modified histone octamer and the 147 base pairs (bp) of DNA wrapped around it^23^. Classically, chromatin has been classified into transcriptionally active, gene-rich euchromatin and silent, gene-poor heterochromatin. However, it is clear that these two broad categories can be further separated based on associated histone modifications, chromatin-binding proteins and function^24–26^. In *Drosophila melanogaster*, the chromatin landscape was classified into five principal states based on the DNA binding profiling of 53 chromatin proteins^27^. Euchromatin was classified into two distinct types: chromatin at highly transcribed, developmentally-regulated genes associated with a SWI/SNF remodeller Brahma (also known as ‘red’) and chromatin enriched in H3K36me3, present at housekeeping loci (‘yellow’)^27^. Heterochromatin was separated into constitutive, H3K9me3-bound regions found mostly at pericentromeres and telomeres (‘green’); facultative, H3K27me3 and Polycomb complex-bound heterochromatin present at developmentally regulated genes (‘blue’); and finally, Lamin-associated regions enriched in histone H1 that lack histone modifications and cover most silent genes (‘black’)^27^. The balance between different types of euchromatin and heterochromatin is regulated by histone modifying enzymes that deposit methylation or acetylation at distinct amino acid residues of histone proteins^28–30^. Histone modifications and chromatin proteins ultimately regulate the spacing of nucleosomes and formation of condensed structures, which affects the accessibility of DNA for transcription^31^.

Here, we investigated the chromatin changes that occur in neural stem cells during quiescence induction and reactivation. We generated a comprehensive genomic dataset of nine chromatin binding proteins and histone modifications in three developmental conditions: proliferating NSCs prior to cell cycle exit, quiescent NSCs and reactivated NSCs. Surprisingly, we found that in quiescent NSCs accessible chromatin is increased at signalling genes as a result of remodelling and deposition of active histone modifications. Strikingly, cell cycle-related loci are retained within permissive, H3K36me3-bound euchromatin but are transcribed at lower levels during quiescence. Overall, our results show that quiescence in NSCs is a process that requires active chromatin remodelling, including increased accessibility and expansion of active chromatin states without, necessarily, concomitant transcription. This may enable quiescent cells to be poised for rapid reactivation.

## Results

### Quiescent neural stem cells gain accessible genomic regions

To investigate chromatin dynamics in neural stem cells *in vivo*, we used Targeted DamID to profile protein DNA binding without cell isolation, immunoprecipitation or fixation^32^. Techniques that rely on cell sorting, such as chromatin immunoprecipitation (ChIP) require high cell numbers per sample and highly specific antibodies^33^. In contrast, Targeted DamID is ideal for profiling genomic binding *in vivo* and when cell sorting can be challenging, such as for small numbers of neural stem cells^32,33^. Targeted DamID is designed to express low-levels of *E.coli* Dam methyltransferase fused to a protein of interest in a tissue-specific manner using the GAL4 system^32^. Dam is guided to specific genomic locations by the protein of interest and methylates the adenine in GATC sequences near its binding sites. This gives rise to methyl groups marking the sites of binding, which can be isolated using methylation-sensitive restriction enzymes. Binding data are normalised against the binding of untethered Dam, which methylates adenines at nucleosome-free GATC regions, yielding cell-specific accessibility profiles similar to ATAC-seq or FAIRE-seq^34^ (Fig. 1A).

**Figure 1.**
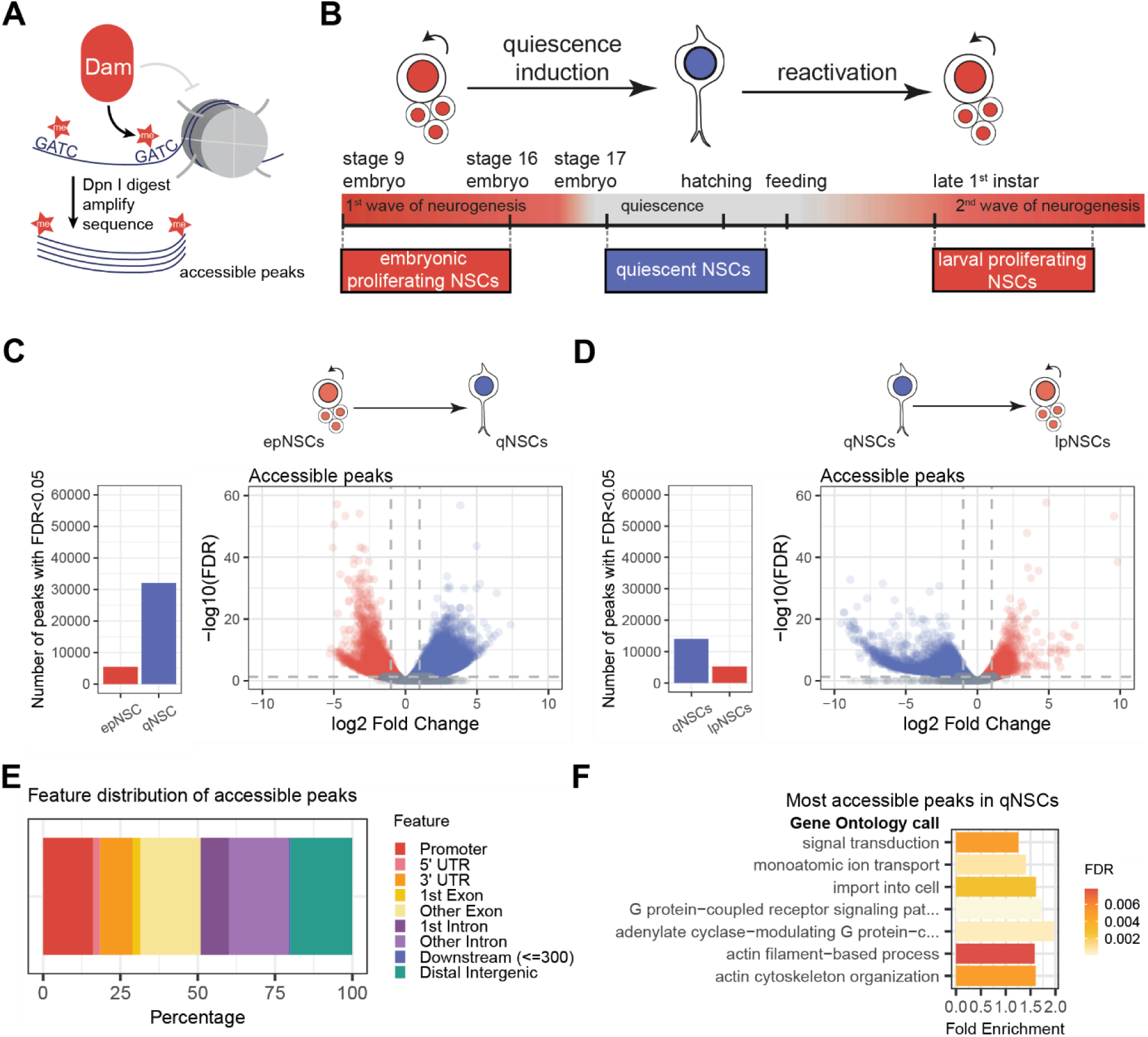
Quiescent neural stem cells gain accessible regions. (**A**) A diagram describing isolation of accessible genomic regions using Targeted DamID. (**B**) Timeline of Dam expression for capture of pre-quiescent (embryonic proliferating, epNSCs), quiescent (qNSCs) and post-quiescent (larval proliferating, lpNSCs). (**B**) Number of significantly differentially accessible peaks (FDR < 0.05) between epNSCs and qNSCs (left) and the volcano showing the fold change and -log_10_ false discovery rate (FDR) for the peaks (right). (**C**) Number of significantly differentially accessible peaks (FDR < 0.05) between qNSCs and lpNSCs (left) and the volcano showing the fold change and -log_10_ false discovery rate (FDR) for the peaks (right). (**E**) Feature annotation of accessible sites gained in quiescent NSCs. (**F**) Gene ontology calls of accessible peaks gained in quiescent NSCs.

To investigate the changes in chromatin accessibility during quiescence and reactivation in NSCs, we drove expression of Dam at different time points with an NSC-specific GAL4 driver (*worniu*-GAL4). First, we profiled NSCs during the embryonic wave of neurogenesis (embryonic proliferating NSCs, epNSCs), from midway through embryogenesis until later embryonic stages (stage 9 through stage 16; according to Campus Ortega and Hartenstein staging^35^; Fig. 1B). The majority of NSCs exit the cell cycle by the end of embryogenesis (stage 17) and begin reactivation during the 1^st^ larval instar^36^. Next, to profile quiescent NSCs (qNSCs), we used a temperature sensitive repressor of the GAL4 protein, GAL80 (GAL80^ts^)^37^ to limit Dam expression to late embryogenesis and early larval first instar (Fig. 1B). Finally, we used GAL80^ts^ to restrict Dam expression to late first instar larvae (24 hours after hatching) to profile larval proliferating NSCs (lpNSCs) (Fig. 1B).

We used differential binding analysis of Dam-enriched fragments to recover differentially accessible genomic peaks. When filtered by false discovery rate (FDR) (FDR<0.01) to retain only significantly different fragments, we observed higher numbers of accessible peaks in quiescent NSCs in comparison to embryonic proliferating or larval proliferating NSCs (Fig. 1C-D). We assayed the genomic distribution of differentially accessible regions and found that chromatin was decondensed upon quiescence induction at both genic and intergenic regions (Fig. 1E). This indicated that qNSCs undergo global expansion of accessible chromatin. This process reverses upon reactivation, indicating that opening of chromatin is a quiescence-specific event (Fig. 1D). Genes with higher number of accessible peaks in qNSCs are associated with Gene Ontology calls related to regulation of the cytoskeleton, cell morphology, and signalling, such as GPCR signalling pathway, actin remodelling and ion transport (Fig. 1F). Therefore, contrary to expectations, we found that quiescent NSCs decondense chromatin at both genic and intergenic regions.

### Neural stem cell chromatin can be classified into seven functional states

Having established that quiescent NSCs gain accessible regions, we investigated the epigenomic changes that enable NSCs to remodel their chromatin through quiescence entry and reactivation. We performed a series of Targeted DamID experiments profiling chromatin modifiers and histone modifications in NSCs. Using the timepoints defined in Figure 1B, we expressed Dam fusion proteins that recognise histone modifications with well-understood functions such as H3K4me3, H3K27me3, H3K9me3, H3K9ac and H4K20me1 (Fig. 2A, Fig. S1, S2). We also expressed Dam fused to chromatin proteins such as the SWI/SNF factor Brahma (Brm), repressive histone H1 and RNA polymerase II (Pol II) (Fig. 2A, Fig. S1, S2).

**Figure 2.**
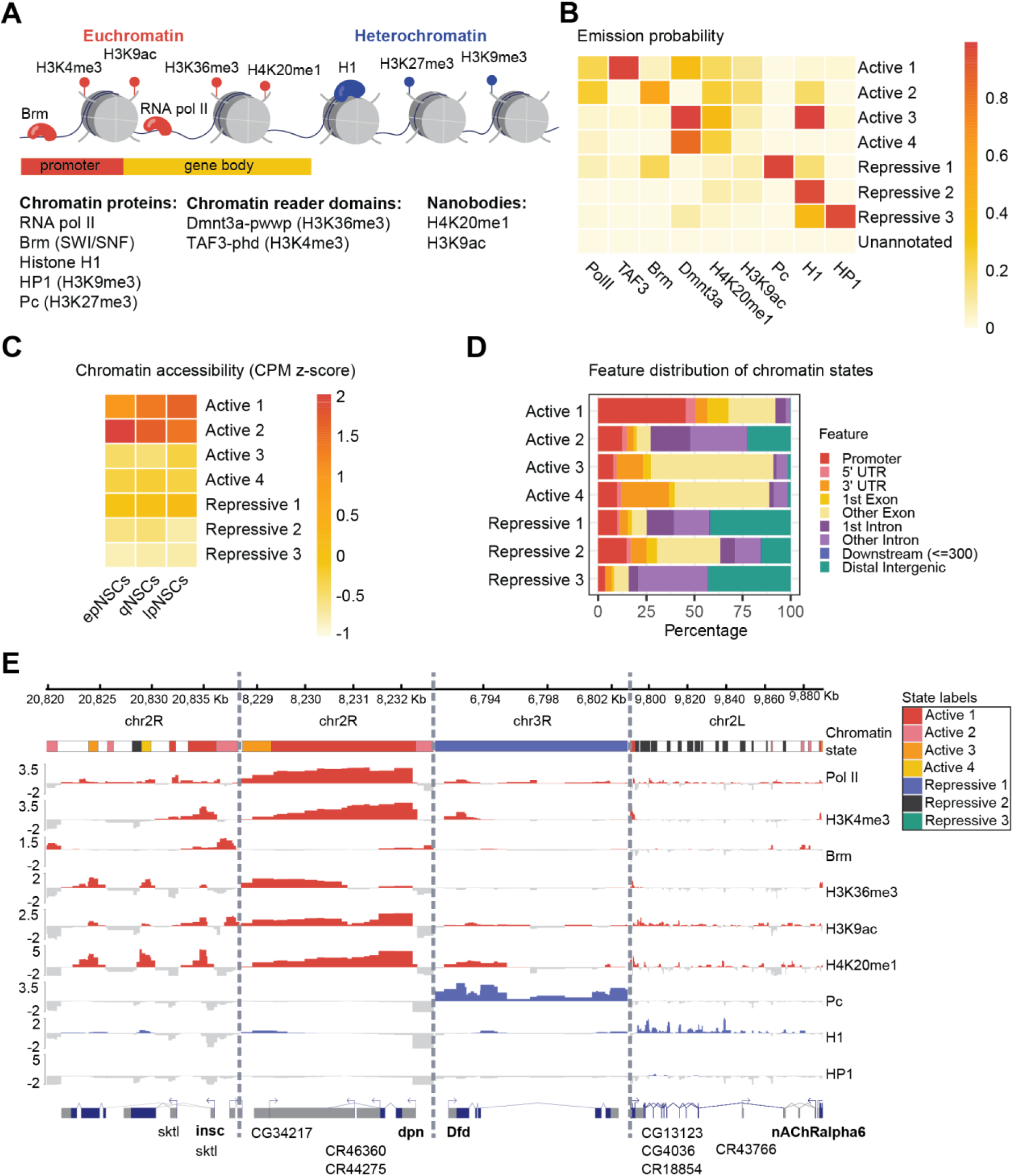
Neural stem cell chromatin can be separated into 7 functional states. (**A**) A diagram showing the repertoire of DamID tools used for hidden Markov modelling of chromatin state data. (**B**) Emission probabilities of the 8-state concatenated chromatin model i.e. how likely is a given signal to be present within a given state. (**D**) Genomic feature distribution of regions annotated as a given chromatin state. (**E**) Genomic tracks of chromatin state annotation and associated Dam fusion signal at four loci in embryonic NSCs. *Inscuteable (insc)* a gene expressed in NSCs, is marked by H3K4me3 at the promoter (Active state 1) and Pol II at the promoter and throughout the gene body together with H3K36me3, H4K20me1 and H3K9ac (Active states 3 and 4). Brahma and H3K9ac are present upstream of the promoter (Active state 2). *Deadpan (dpn)* is a highly expressed NSC marker. As such, it is mostly annotated by state 1, with high levels of active histone modifications and Pol II throughout the locus. *Deformed (Dfd)* is a Hox gene repressed by Polycomb in epNSCs and labelled as Repressive state 1 accordingly. Finally, acetylcholine nicotinic receptor 6 alpha (nAChRalpha6) is not expressed in proliferating NSCs and is mostly marked by repressive histone H1 and labelled as Repressive state 2.

We used machine learning to create chromatin state annotations based on the genomic binding profiles of these nine Dam fusions. We fitted a concatenated hidden Markov model for the three neural stem cell conditions to our datasets with ChromHMM^9^ (explained in detail in Fig. S3A,B). We identified four active and three repressive chromatin states in the 8-state model (Fig. 2B,C). The model classified chromatin in accordance with those previously obtained by state modelling in different *Drosophila* tissues and cells^27,38–40^. To maintain continuity with previous annotations, we used a similar colour labelling code to denote different chromatin states (Fig. 2E).

Active state 1 (red) is characterised by high levels of H3K4me3, Pol II and other active histone modifications, and is found mostly around the transcriptional start sites (TSSs) and at short, highly expressed genes, such as the NSC gene, Deadpan (Fig. 2B-E). Active state 2 (pink) is characterised by high levels of Brm, Pol II and other active chromatin marks. Brm is a *Drosophila* ATPase dependent chromatin remodeller, homologous to SWI/SNF factor^41^, that can modulate H3K27ac via its association with histone acetyltransferase CBP/nej and H3K27me3 demethylase Utx^10^. Many regions annotated as Active state 2 are found at introns and distal intergenic regions (Fig. 2D,E). Chromatin state associated with H3K27ac, and present in non-coding or intronic regions, suggests that it marks active enhancers^42^.

Active states 3 and 4 (orange and yellow) exhibit characteristics of transcriptional elongation at gene bodies, such as H3K36me3 and H4K20me1^11,12^ (Fig. 2B,E). The identity of Active state 3 and 4 is supported by their presence in exons (Fig. 2D,E). Active state 1 is often followed by Active state 4 in the genome, reflecting their localisation at promoters and active gene bodies, respectively (Fig. 2E,S3C). Importantly, chromatin accessibility is highest in states 1 and 2, at the regions associated with regulatory elements (Fig. 2C). Previous chromatin state annotations in *Drosophila* have separated two types of euchromatin: Trithorax-associated ‘red’ regions and H3K36me3-associated ‘yellow’ regions^27,38,39,42^. In our model, Active states 1 and 2 are most closely related to ‘red’ chromatin, and Active state 3 and 4 are related to ‘yellow’ chromatin.

The remaining states are characterised by lower chromatin accessibility and the presence of chromatin proteins that suggest repressive chromatin (Fig. 2B,C). We observe a clear separation of Polycomb-bound regions (‘blue’ chromatin in previous studies^27,38,39^ and here) and HP1-related heterochromatin (‘green’ in previous studies^27,38,39^ and here), representing Repressive states 1 and 3, respectively (Fig. 2B,C,E, S3C). Repressive state 2 is reminiscent of the ‘black’ chromatin reported previously in *Drosophila*^27,38,39,42^ or ‘naïve’ chromatin in mammalian cell lines^24^, and consists of repressive regions with fewer histone modifications and high histone H1 levels (Fig. 2B). Finally, the regions that lacked signal, including histone H1, were named ‘Unannotated’ and excluded from most further analyses (Fig. 2B).

### Quiescent NSCs gain active chromatin domains

Having assigned identities to the chromatin states, we assayed the genomic frequency of each state in NSC genomes during embryonic proliferation (epNSCs), quiescence (qNSCs) and reactivation (lpNSCs). Remarkably, we found that the fraction of Active states 1 and 2 (red and pink) increased in quiescence, confirming the results we obtained investigating chromatin accessibility alone (Fig. 3A). The percentage of Active state 1 decreased during reactivation, suggesting that this chromatin transition is specific to quiescence (Fig. 3A). To confirm that the increase of Active state 1 is specifically associated with opening of promoters, we compared the chromatin state frequency at TSS-associated regions (Fig. 3B). We defined the putative *Drosophila* promoter region as 500bp upstream to 200bp downstream of the TSS and annotated each TSS region with a chromatin state. We found that Active state 1 increase in quiescence is more pronounced at TSS-associated regions. To validate this finding using analysis independent of the chromatin state model, we assayed the number of H3K4me3-bound peaks during quiescence induction with differential binding analysis and found, accordingly, that the number of H3K4me3-bound regions was higher in qNSCs (Fig. 3C). Overall, we showed chromatin remodelling that leads to higher number of accessible regions in quiescent NSCs is due to gain of Active states 1 and 2 associated with promoter and enhancer regions. This result was striking, as chromatin was shown to condense and gain repressive histone methylation in other models of quiescence^9^.

**Figure 3.**
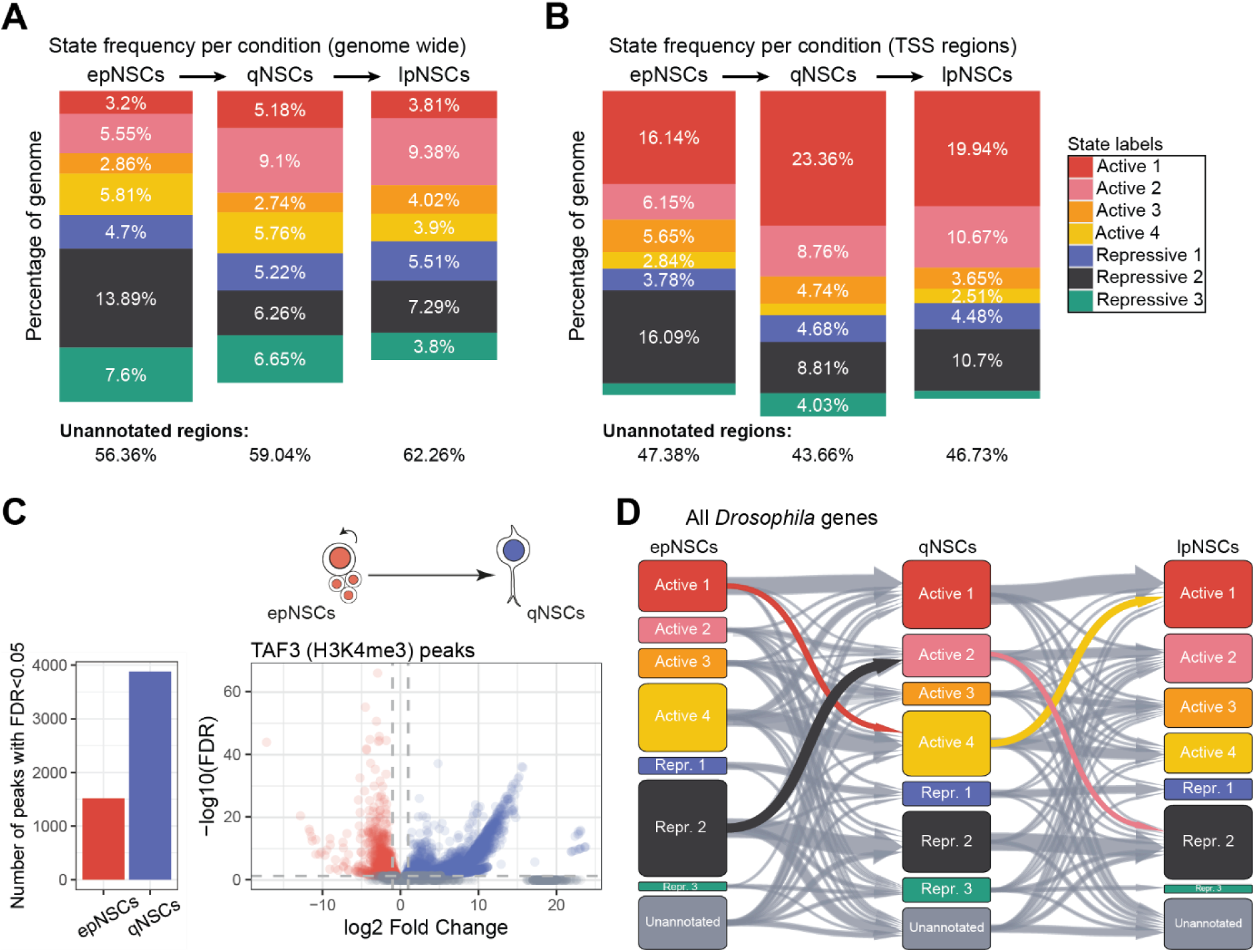
Quiescent NSCs gain domains marked by active chromatin states. **(A)** Chromatin state frequency at the whole genome in the 3 tested conditions. (**B**) Chromatin state frequency at state-annotated promoters (defined as regions 500bp upstream and 200bp downstream of the TSS) in the 3 tested conditions. (**C**) Number of significantly differentially bound H3K4me3 (TAF3) peaks (FDR < 0.05) between epNSCs and qNSCs (left) and the volcano showing the fold change and -log_10_ false discovery rate (FDR) for the peaks (right). (**D**) Alluvial plot for gene-based chromatin state transitions for all *Drosophila* genes. Genes which remain in unannotated chromatin in all 3 conditions are filtered out. Transitions assayed in later figures are highlighted.

### Quiescent NSCs maintain permissive chromatin at cell cycle genes

We were surprised to find increased open chromatin during quiescence, given that chromatin accessibility is generally thought to represent areas of active transcription. We asked if specific chromatin state transitions were associated with proliferation genes and with genes that are transcribed in quiescent NSCs. To better reflect chromatin transitions at the single gene level, we developed a new bioinformatic tool (*feat_annot*) to annotate each *Drosophila* locus with a chromatin state (Fig. S3D). Although a single gene is likely marked by multiple states, we used the chromatin state with the highest coverage at given loci for gene annotation to reduce dimensionality of the data. We used this annotation to plot the frequency of chromatin state transitions at all *Drosophila* genes and found unique chromatin transitions that occur between active embryonic and quiescent NSCs both within euchromatin and heterochromatin (Fig. 3D).

We focused on a group of 325 genes that transitioned from Active state 1 (equivalent to ‘red’ euchromatin according to classification by Filion et al.^27^) to H3K36me3-associated Active state 4 (‘yellow’ euchromatin) state during quiescence induction (Fig. 4A, B). As Active state 4 (yellow) is associated with lower Pol II levels and chromatin accessibility (Fig. 2B,C), we speculated this gene group would be downregulated during quiescence. Strikingly, we found that most genes within this category are necessary for the progression of the S and M phases of the cell cycle based on Gene Ontology calls, including DNA replication, organisation of the mitotic spindle and chromatid separation (Fig. 4C). State 3 and 4 have lower chromatin accessibility and Pol II levels than state 1 (Fig. 2C), which indicates that these loci are within permissive, but not highly active chromatin state.

**Figure 4.**
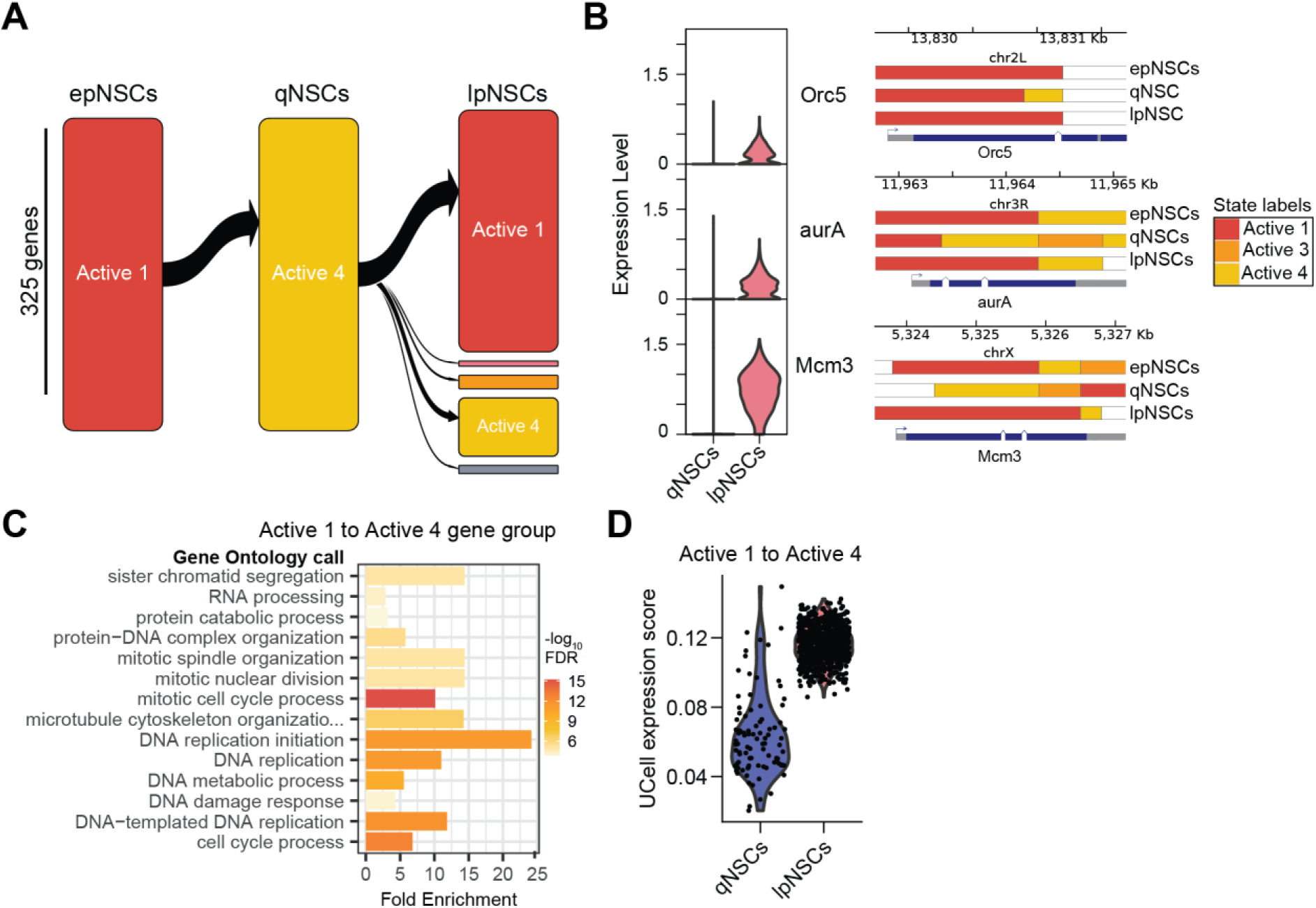
Quiescent NSCs maintain euchromatin at cell cycle genes. (**A**) Alluvial plot for genes undergoing a transition from active state 1 to state 4 during quiescence induction (325 genes). (**B**) mRNA expression levels and chromatin state annotation at three example loci: *Origin recognition complex subunit 5* (*Orc5*), *aurora A* (*aurA*) and *Minichromosome maintenance complex component 3* (*Mcm3*). (**C**) Gene ontology (Biological Process Slim) of genes shown in (**A**). (**D**) UCell expression score of gene set in (**A**). All gene expression results are based on scRNA-seq dataset from ref.^43^.

To assess transcription levels at these loci, we analysed our single cell RNA sequencing dataset of the embryonic and larval brain^43^. Several genes necessary for proliferation (*aurora A, Orc5* and *Mcm3* shown) were maintained in active states in quiescent NSC (Active state 1, 3 and 4; red, orange and yellow), but were not transcribed, as indicated by single cell RNA-sequencing of quiescent and larval proliferating NSCs performed previously by our group^43^ (Fig. 4B). Finally, we calculated the expression score (UCell score^44^) for genes moving between Active state 1 and Active state 4 in quiescent and larval proliferating NSCs. We found that cell cycle genes that transition from Active state 4 to Active state 1 during reactivation increase their expression only in lpNSCs (Figure 4D). This indicates that genes required for M and S phase of the cell cycle are within permissive chromatin during quiescence but not highly transcribed.

### Quiescent NSCs derepress signalling genes

The transition of a large group of genes from repressive H1 chromatin (Repressive 2) in embryonic proliferating NSCs to active euchromatin associated with Brm (Active 2) in quiescent NSCs suggested a group of genes that is specifically de-repressed during quiescence (Fig. 3D). We assayed the putative ‘quiescence expressed’ group of loci characterised by transition out of Repressive state 2 (black) to Active state 2 (pink) (Fig. 5A). This group consisted of over 700 genes with ‘neuronal-like’ functions such as synaptic and neurotransmitter signalling, ion transport across membranes, cell adhesion and microtubule remodelling based on Gene Ontology calls (Fig. 5B). We found that many of these genes transition back to Repressive state 2 (black) in lpNSCs, indicating they become repressed during reactivation (Fig. 5A). Importantly, this suggests that transcription of neuronal genes only occurs in quiescent NSCs and is not a feature of neural stem cells in general.

**Figure 5.**
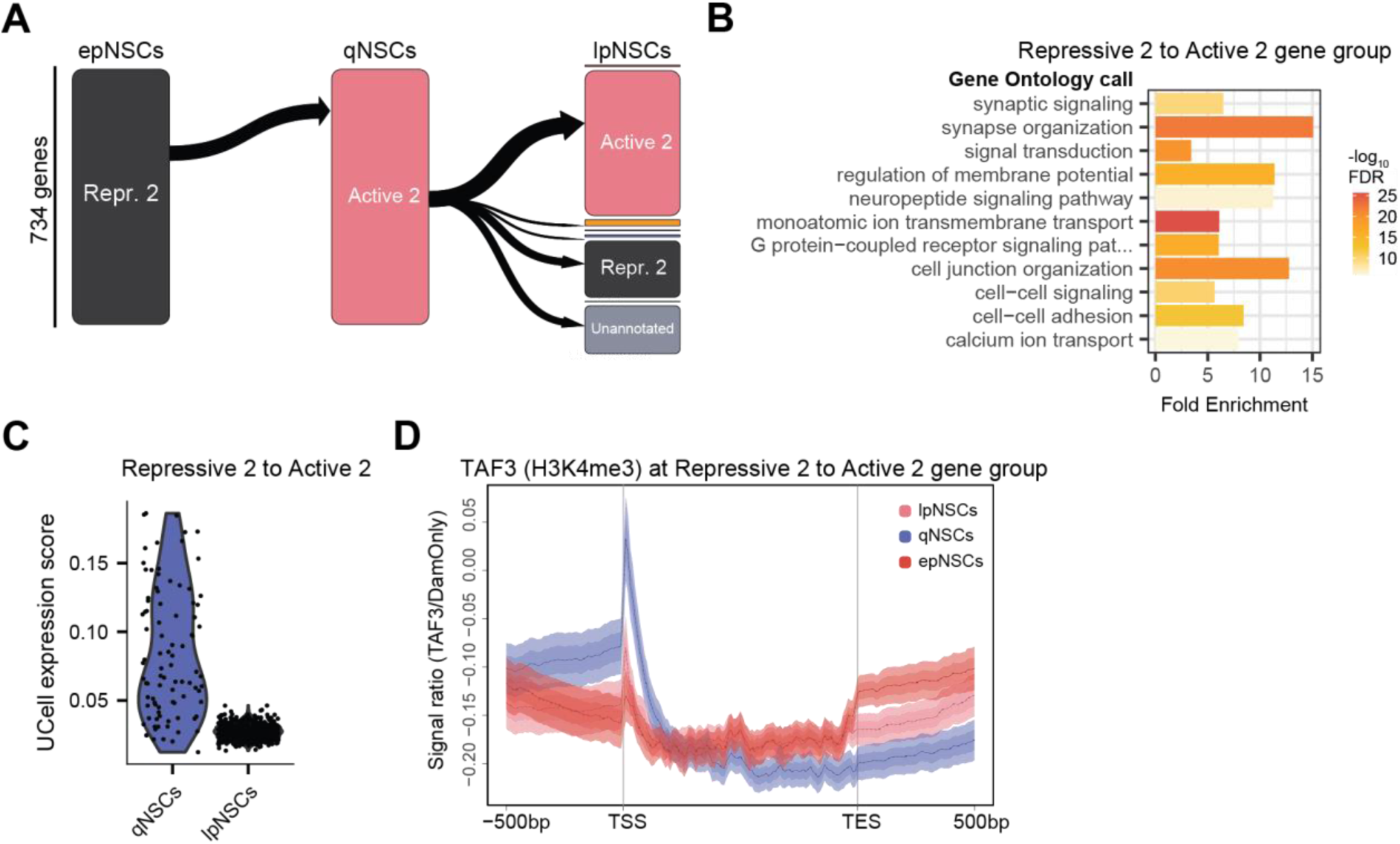
Quiescent NSCs derepress neuronal signalling genes. (**A**) Alluvial plot for genes undergoing a transition from repressive state 6 to active state 2 during quiescence induction (734 genes). (**B**) Gene ontology (Biological Process Slim) of genes shown in (**A**). (**C**) UCell expression score of gene set in (**A**). All gene expression results are based on scRNA-seq dataset from ref.^43^. (**D**) Signal ratio of the TAF3-Dam fusion (reflecting H3K4me3) at the gene group shown in (**A**) in embryonic proliferating (epNSCs), quiescent (qNSCs) and larval proliferating NSCs (lpNSCs).

We validated expression of genes undergoing Repressive state 2 to Active state 2 transition in quiescent (qNSCs) and reactivated (lpNSCs) neural stem cells using scRNA-seq^43^ and found that these genes are transcribed in quiescent cells, but not in reactivated cells (Fig. 5C). Accordingly, we have assayed the presence of H3K4me3 at this group of genes and found that the peak of H3K4me3 signal is highly enriched at the TSS in quiescent NSCs in comparison to embryonic proliferating or reactivated NSCs (Fig. 5D). Interestingly, quiescent NSCs change morphology during quiescence induction from large, round cells to small cells that extend a basal process towards the neuropile of the larval CNS^45^. This suggests that genes with functions related to building the basal process and cell signalling, including microtubule binding proteins, ion channels, neurotransmitter receptors and synaptic proteins are briefly derepressed during quiescence but cease transcription upon reactivation, in accordance with our previous results^43^. In fact, we also previously showed that over 48 hours after reactivation, genes with ‘signalling’ and ‘neuronal’ functions are still repressed by H1 ‘black’ chromatin^39^ (here named Repressive 2). In conclusion, we found that chromatin remodelling in quiescent NSCs enables them to undergo reversible cell cycle arrest whilst inducing expression of genes necessary for signalling and extension of the basal projection.

## Discussion

The balance between neural stem cell proliferation and quiescence, as well as the proper order of reactivation events, is crucial to maintain tissue homeostasis. Previous studies have shown that quiescent cells are characterised by accumulation of heterochromatin, low metabolic rate and low transcription levels^10,46–49^. Here, we show that one of the main chromatin transitions in quiescent neural stem cells is an increase in chromatin accessibility and active chromatin states (Fig. 6A). This remodelling is the result of transition from H1 bound chromatin (Repressive state 2) to a Brm-marked state (Active state 2) during quiescence induction. For instance, genes necessary for axonal fasciculation (*beaten path IIIa, beat-IIIa*) and calcium signalling (*Ca^2+^-channel protein α_1_ subunit D, Ca-alpha1D)* move from Repressive state 2 to active state 1 in quiescent NSCs. By late first instar, some neuronal genes have moved back to Repressive state 2 (Fig 6B). During reactivation, neural stem cells retract the basal process, increase their size and adopt a rounded shape. Therefore, downregulation of microtubule binding and synaptic genes during reactivation could be necessary for disassembly of the quiescent neural stem cell projection. We previously reported that repressive H1 ‘black’ chromatin marks neuronal genes in proliferating NSCs in third instar larvae^39^, indicating that the brief de-repression of neuronal genes is specific to quiescence. At the same time, genes necessary for cell cycle progression, such as the *polo kinase* (*polo*) and Rho GTPase *tumbleweed* (*tum*) move from Active state 1 chromatin to Active state 4 permissive euchromatin marked by H3K36me3 (Fig 6C). This transition is associated with reduced transcription and is reversed upon reactivation, when NSCs start transcribing genes needed for replication and mitosis.

**Figure 6.**
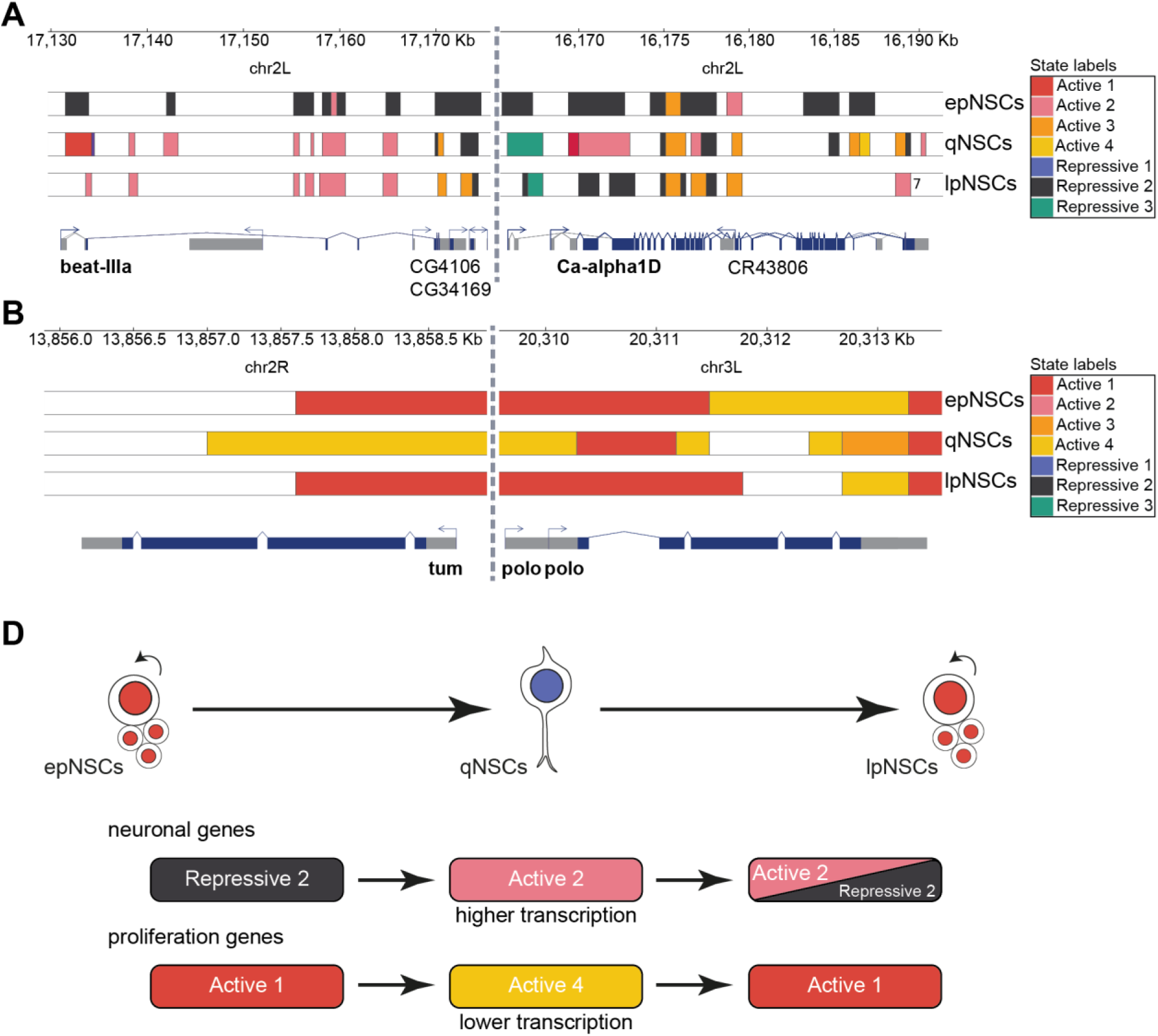
Quiescent NSCs undergo expansive chromatin remodelling. (**A**) Example genomic tracks of two neuronal genes that undergo the Repressive state 2 to Active state 2 transition during quiescence induction: *beaten path IIIa* (*beat-IIIa*) and *Ca^2+^-channel protein α_1_ subunit D (Ca-alpha1D).* (**B**) Example genomic tracks of two proliferating genes that undergo the Active state 1 to Active state 4 transition during quiescence induction: *tumbleweed* (*tum*) and *polo kinase* (*polo*). (**C**) Model of chromatin remodelling in NSCs. During quiescence induction, NSCs exhibit increased chromatin accessibility due to de-repression of neuronal genes which transition from state 6 (H1-bound) chromatin to active (Brahma-bound) euchromatin associated with higher transcription. As NSCs reactivate, they begin to slowly transition back to H1-bound Repressive state 2 chromatin and lower their transcription levels, although some genes have transitioned back to repressive chromatin by late first instar. At the same time, genes necessary for proliferation (specifically M and S phase) transition from H3K4me3-marked Active state 1 associated with high expression levels to permissive euchromatin characterised by H3K36me3 (Active state 4) and lower transcription. As NSCs reactivate, proliferation genes become upregulated due to a transition back to Active state 1.

Quiescent stem cells re-enter the cell cycle in response to specific stimuli. Reactivation can occur within hours, such as in response to dietary signals in *Drosophila* NSCs. To ensure a rapid response, it might be most effective for proliferation genes to remain within permissive chromatin and boost their transcription once NSCs are reactivated. A similar process has been reported in quiescent myoblasts, where the *cyclinA2* locus is found in permissive chromatin, marked by both H3K4me3 and H3K9me2^50^, but not transcribed. Permissive euchromatin at *cyclinA2* inhibited deposition of H3K27me3, and therefore hyper-repression of the locus that would block or slow down cell cycle re-entry upon reactivation^50^.

Quiescent NSCs also reactivate within a specific sequence to maintain the progenitor pool and build appropriate neural circuits. We previously found that *Drosophila* quiescent NSCs transiently transcribe neuronal genes that encode synapse-associated proteins and ion channels needed for depolarisation^43^. Disruption of membrane polarisation led to an abnormal order of NSC reactivation^43^. Here, we have shown that genes identified previously by single cell RNA sequencing^43^ are part of a larger group of loci that undergo a chromatin transition from H1 histone-marked Repressive state 2 (black) heterochromatin to ‘Active state 2’ bound by Brm (pink). These ‘neuronal signalling’ genes are found in H1-bound repressive chromatin in embryonic proliferating NSCs, and quiescent NSCs briefly derepress those loci.

The mobilisation of nucleosomes and eviction of histone H1, which leads to an increase in chromatin accessibility, can be induced by sequence-specific pioneering transcription factors that bind to nucleosomes^51^. Crucially, this can occur prior to SWI/SNF recruitment to the locus^51^. We hypothesise that sequence-specific transcription factors recognising motifs present in promoters of the genes that become de-repressed in qNSCs (e.g. the GAGA factor, GAF) could potentially induce the Repressive state 2 to Active state 2 transition (Fig. S4A-B). We would therefore anticipate that disruption of chromatin remodellers, or sequence specific transcription factors, could impact the proper order of reactivation of NSCs by preventing expression of the signalling machinery.

The evidence for specific chromatin effectors in regulation of NSC function is limited. For instance, loss of an ATP-dependent chromatin remodeller, Chd7, in the mouse subgranular zone of the hippocampus increased the number of cycling cells, depleting the stem cell pool over time^52^. Recent studies of mammalian NSC chromatin attributed loss of functionality of quiescent NSCs with age in mouse to decreased chromatin accessibility, leading to downregulation of cell adhesion genes and ultimately, loss of adherence of quiescence cells^14^.

In an *in vitro* neural stem cell progenitor (NSPC) quiescent system, activated NSPCs were shown to gain accessibility at H3K27ac-marked enhancers found in distal intergenic regions, whereas promoters remained stably accessible during reactivation^15^. Finally, BMP-4 induced quiescence in mouse NSCs *in vitro* led to activation of over 6000 enhancers, twice as many as the number of active enhancers in proliferating NSCs^53^, indicating that it is likely that chromatin accessibility also increases in mammalian NSCs.

Chromatin accessibility has been associated with quiescence in the context of B cell lymphocytes^11^, whereas decreased histone methylation (both active and repressive) characterises hair follicle stem cells^13^. As pluripotency is associated with open chromatin^54^, this mechanism may enable NSCs to retain their plasticity and ability to respond to different environmental stimuli. In conclusion, we found that in contrast to other quiescence systems, in which cells condense chromatin and shut down nuclear processes (such as in nutrition-deprived yeast^6^), neural stem cells actively remodel their chromatin to induce functional expression of signalling components and safeguard proliferation genes against hyper-repression.

## Methods

### Fly Stocks

The following fly stocks were used to drive expression of transgenes: *worniu-*GAL4^55^, *tub*-GAL80^ts^ (BDSC 7019)*, UAS-LT3-NDam* and *UAS-LT3-NDam-RNAPol II*^32^*, UAS-LT3-Dam-Pc, UAS-LT3-Dam-HP1a, UAS-LT3-Dam-Brm, UAS-LT3-Dam-H1*^39^*, UAS-LT3-Dam-TAF3, UAS-LT3-Dam-Dmnt3a, UAS-LT3-Dam-15F11, UAS-LT3-Dam-19E5*^56^.

### Targeted DamID experimental design

To perform Targeted DamID, *wor*-GAL4 or *wor*-GAL4, tubGAL80^ts^, depending on the condition were crossed to flies carrying a Dam fusion. UAS-LT3-Dam was used as a control for all experiments and three or more replicates were performed for each condition.

For the embryonic proliferating NSC (epNSC) condition, Dam fusion lines were crossed to *wor*-GAL4. Flies were reared in cages at 25°C and allowed to lay eggs on apple juice plates for 1 hour. The plates were kept at 25°C for 12h and 50µl of embryo pellet per replicate was collected into microcentrifuge tubes and frozen at -70°C.

For the quiescent NSC (qNSC) condition, Dam fusion lines were crossed to *wor*-GAL4, tubGAL80^ts^. Flies laid eggs on apple juice plates for 1 hour and the plates were then transferred to 18°C for 28h to prevent expression of Dam with the use of the temperature-sensitive GAL80 repressor, restricting the transgene expression to after embryonic stage 17. To induce expression of Dam, the plates were shifted to 29°C for 12h. About 300 whole larvae per replicate were collected into microcentrifuge tubes and frozen at -70°C.

For the larval proliferating NSC (lpNSC) condition, Dam fusion lines were also crossed to *wor*-GAL4, tubGAL80^ts^. Flies laid eggs on apple juice plates for 1 hour and the plates were then transferred to 18°C for 56h, restricting transgene expression to late first larval instar (equivalent to 24h after larval hatching, ALH, at 25°C). The plates were then shifted to 29°C for 12h. About 200 whole larvae per replicate were collected into microcentrifuge tubes and frozen at -70°C.

### Targeted DamID library processing

Targeted DamID were processed according to a published DamID-seq protocol^57^. Briefly, genomic DNA was extracted with the QIAamp DNA Micro Kit and digested with DpnI and DpnII to isolated methylated DNA. The fragments are PCR amplified and sonicated before generating a sequencing library and multiplexing with a homebrew Truseq kit. Sequencing was performed as single end 50 or 100 bp reads generated by an Illumina HiSeq 1500 or Illumina NovaSeq 6000, dependent on the dataset, at the Gurdon Institute NGS Core Facility.

### Targeted DamID data processing

Fastq files were analysed with damidseq_pipeline (vR.1)^58^, using the damMer suite for parallelisation and pairwise comparisons of DamID replicates^57^. Briefly, damidseq_pipeline was used to align fastq files with bowtie2^59^ and extend the reads towards 300bp or the first GATC site. Next, the bam files were used as input to generate normalized binding tracks in bedgraph format. DamMer was used to parallelise each Dam-Only to Dam-fusion comparison and perform quantile normalisation between comparisons and averaging. The variability of the extended *.bam files was visualised using Pearson correlation performed using deeptools multiBamSummary and plotHeatmap-pearson commands. Extended reads were also used to perform MACS2 broad peak calling for all pairwise comparisons between Dam-fusion and Dam-only replicates. Peaks were thresholded based on FDR<10^-5^, merged and filtered based on appearance in more than 50% replicates. Peaks were also called on Dam-only replicates alone, serving as a chromatin accessibility readout.

### Generating hidden Markov models

Peak files were used as input to generate hidden Markov model using ChromHMM^60^. Reproducible peak files were binarised with BinarizeBed with bin size of 300bp and the ‘peaks’ option in a concatenated format (each NSCs condition used as input to learn one model). Learn Model command was executed for each condition with a differing number of states between 5 and 12. The resulting emission probabilities of 5-12-state models were compared and the 8-state models was selected as the most relevant for further analyses.

### Examination of chromatin states

Feature annotation of chromatin states bins was performed with ChiPSeeker^61^. Visualisation of emission and transition probabilities derived from ChromHMM and chromatin state frequency was performed using custom R scripts. Dam-Only read coverage per chromatin state was calculated with a custom script (*coverage_over.sh*), which used bedtools coverage to calculate read count per million (CPM) per state and visualised it using pheatmap R package.

### Chromatin state annotation

A custom script was created (*feat_annot.sh*) to annotate genomic features (genes and TSS regions) with the chromatin state that is most prevalent within that DNA region. Briefly, genome annotation files generated by ChromHMM were split into separate files based on the chromatin state. Bedmap was used to calculate the base percentage of a chromatin state at given feature. For the TSS regions, a region 500bp upstream and 200bp downstream was used as reference. Chromatin state with the highest base percentage was treated as the state annotation for given feature. The ‘unannotated’ state was thresholded at 75% i.e. the ‘unannotated’ state was only called when a feature had a base percentage higher than 75%, otherwise the second most prevalent state was called. These annotations were used to plot chromatin state frequency at the TSS and to perform all alluvial plots, which were generated with *Transition*-class from the ‘Gmisc’ R package.

### Differential binding analysis

For comparison of chromatin accessibility between NSC conditions, the DiffBind R package was used^62^. Broadpeak and extended read files of the Dam-only dataset were used as input for differential accessibility analysis. For differential binding of H3K4me3 peaks, TAF3 broadpeaks and bam files were used as input, with the Dam-only bam files as control. Default settings were used for majority of commands, apart from dba.count, where bSubControl=TRUE, minOverlap=1, fragmentSize = 0, summits = TRUE.

### Gene ontology

Gene ontology was performed with PANTHER18.0^63^ as a statistical overrepresentation test (Fisher’s Exact text with False Discovery Rate correction) of the GO biological process complete set, using whole Drosophila gene set as reference. The database was accessed used rbioapi package in R and the results were visualised with custom R scripts.

### Single cell RNA-sequencing dataset analysis

A previously published scRNA-seq dataset of NSCs at embryonic stage 17 and 24h ALH^43^ was analysed using the Seurat suite (v4)^64^. Cells in the two datasets were filtered based on RNA count and mitcQC, merged and sctransform normalization was applied. The two datasets were integrated and clustered. NSC-specific cluster that expressed dpn and other NSC-markers was subsetted and the dataset was normalised and scaled within that cluster. Gene group expression score was calculated using UCell^44^.

### Motif analysis

Putative promoter regions of the ‘Repressive state 2’ to ‘Active state 2’ gene group were defined as 500bp upstream and 200bp downstream of the TSS and recovered from the dm6 genome. Motif discovery and comparison to known motif databases was performed with Homer (v5.1) using the findMotifsGenome.pl command with the -size given option.

### Data visualisation

All graphs were generated using ggplot2 (v3.4.2) or pheatmap (v. 1.0.12) unless stated otherwise. Figures were assembled using Adobe Illustrator (v27.5).

### Materials availability

Plasmids and fly stocks used in this study are available upon request.

## Data and code availability

Targeted DamID datasets are available upon request. All datasets will be deposited at GEO and will be publicly available as of the date of publication. Single cell RNA sequencing data has been previously reported at ref^43^. All original code and details of bioinformatic analyses will be publicly available as of the date of publication at Github.

## Acknowledgements

We thank the Gurdon Institute Sequencing Facility for sequencing of DamID samples. We would like to thank Adam Reid for his advice on differential binding analysis. This work was funded by Wellcome Trust Senior Investigator Award (103792), Wellcome Investigator Award (223111) and Royal Society Darwin Trust Research Professorship to A.H.B (RP150061). A.M. was supported by a Wellcome Trust PhD Studentship (222276/Z/20/Z). A.H.B acknowledges core funding to the Gurdon Institute from the Wellcome Trust (092096) and CRUK (C6946/A14492).

## Author contributions

A.M., J.A. and A.H.B conceptualised the project. A.M and J.A. generated DamID datasets.

A.M. performed all bioinformatics analysis. A.M. and A.H.B wrote the manuscript.

