## Supplementary Figures for "Neural stem cell quiescence is actively maintained by the epigenome"

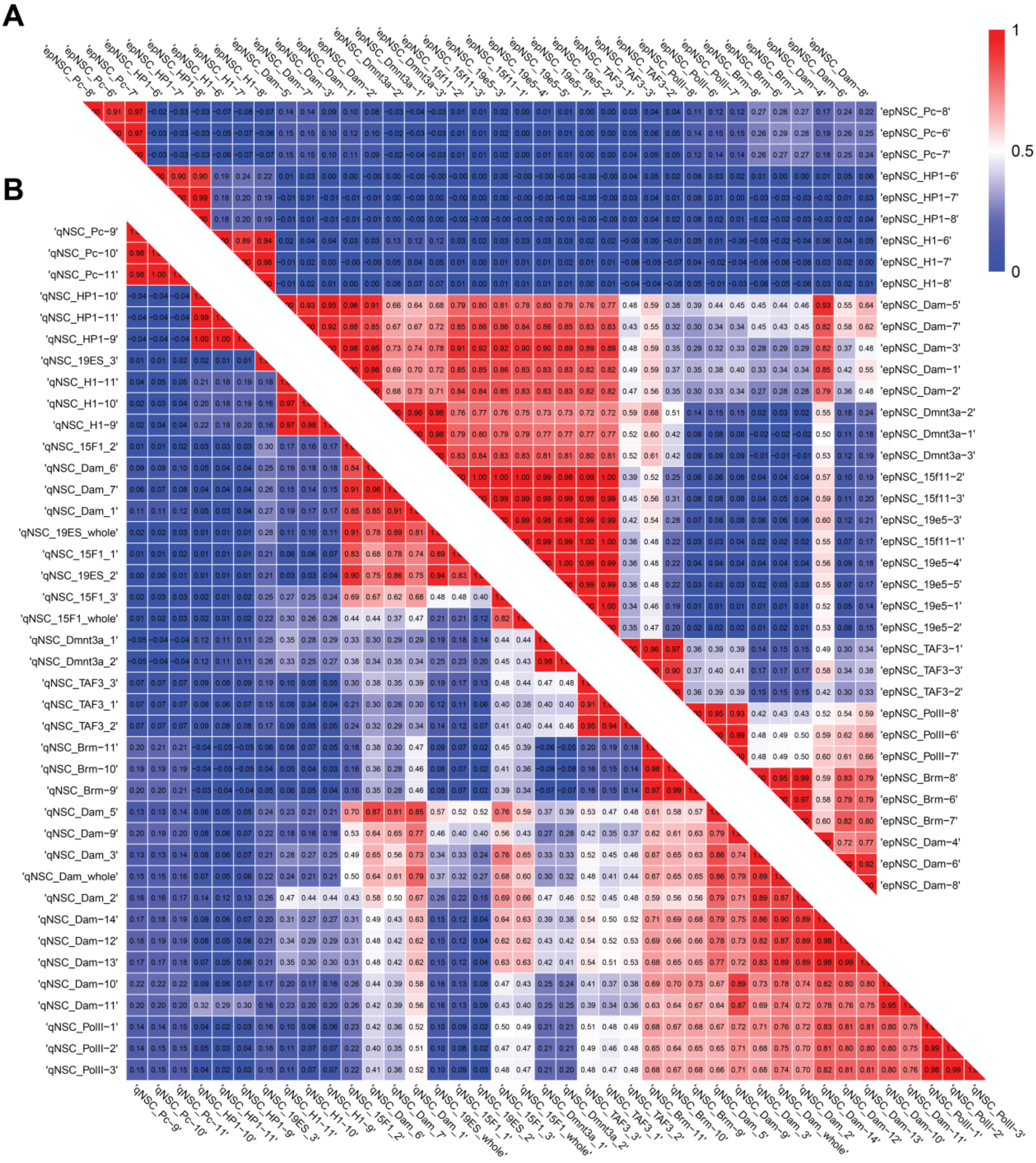


**Supplementary Figure 1. Heatmap of Pearson correlation of DamID read libraries (bam files).** (**A**) Correlation of epNSCs replicates. (B) Correlation of qNSCs replicates.

**A**

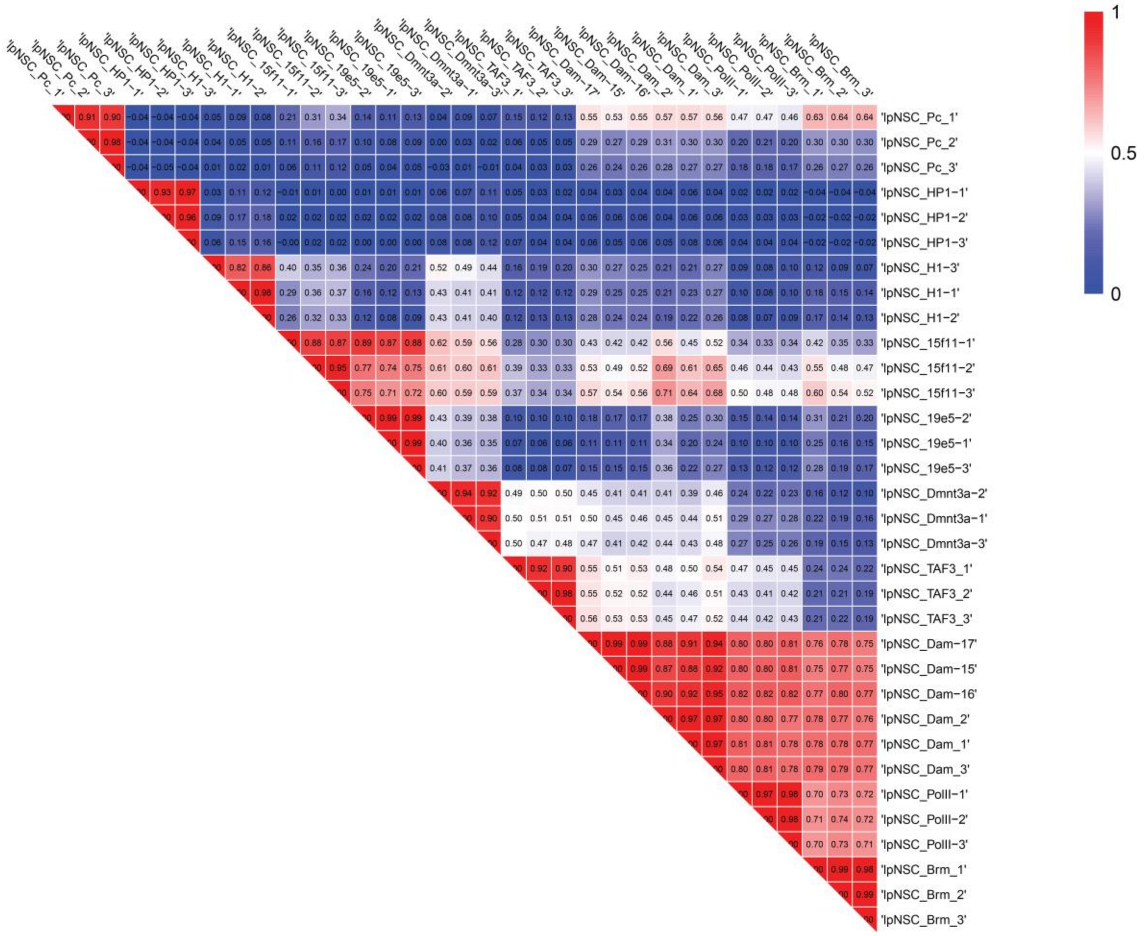


**Supplementary Figure 2. Heatmap of Pearson correlation of lpNSCs DamID read libraries (bam files).**

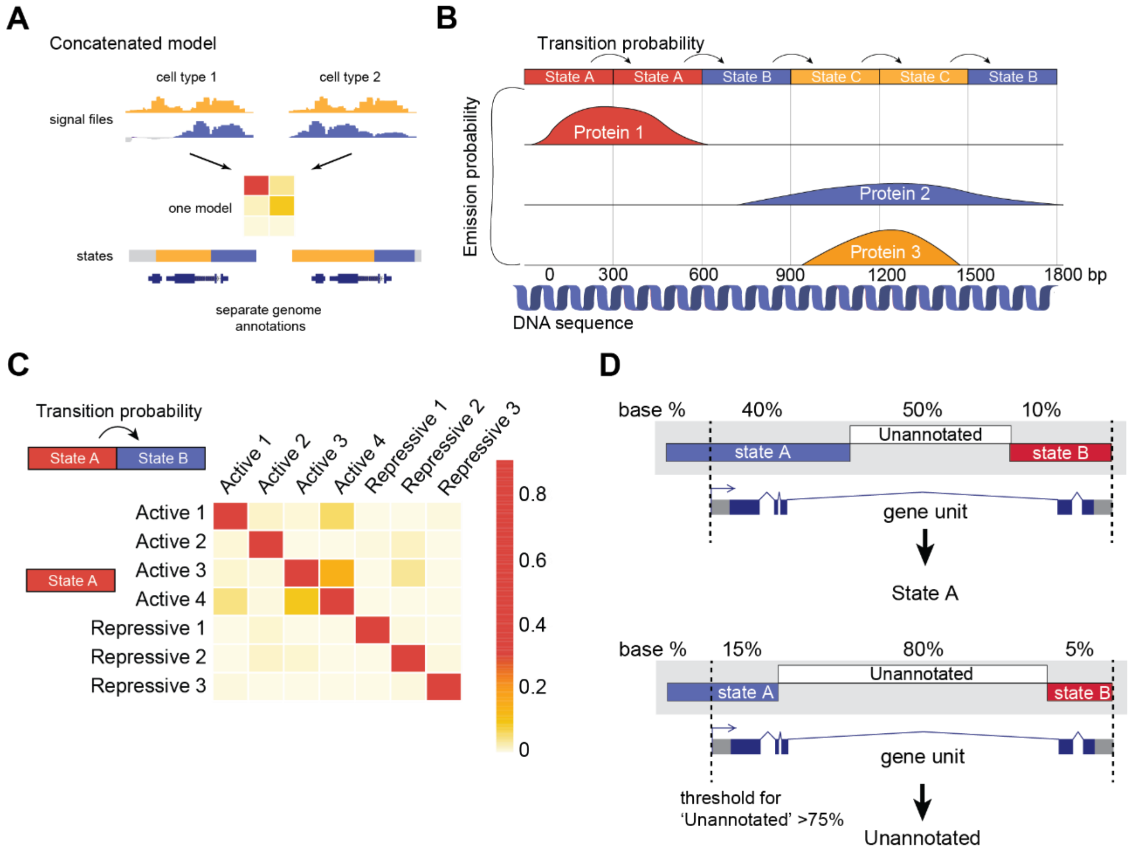


**Supplementary Figure 3. Examination of chromatin state model.** (**A**) A diagram of a concatenated chromatin state model. Concatenated model is learned based on input from multiple conditions (cell populations). Single matrices of emission probabilities and transition probabilities for all conditions is calculated, however, genomic annotations are generated based on model separately for each condition. (**B**) An explanation of emission and transition probabilities of a hidden Markov model (HMM). To generate a chromatin state HMM, the genome is binned (here in equally sized bins of 300bp). The emission probability is a likelihood of given signal (here Targeted DamID fusion signal) being present at the bin labelled as a given state. The transition probability is a likelihood of a given state (label) being followed by a different state in a linear sequence (here the genetic sequence). Importantly, since Targeted DamID signal does not carry any strand information, the transition probability cannot differentiate the strandedness of a genetic locus, i.e. state characteristic of promoters can either follow or be followed by a state characteristic of the gene body, dependent on which strand the gene is encoded. (**C**) Transition probabilities of the 8-state concatenated chromatin model i.e. how likely is one state to follow another. (**D**) Diagram for gene annotation with chromatin states by *feat_annot.sh* script. First, the percentage of base pairs covered by a given chromatin state in each locus is calculated. The chromatin state with the highest base pair coverage is used to annotate the associated locus. As longer genes are likely to be predominantly labelled as the 'Unannotated' state, a locus is only labelled as 'Unannotated' if over 75% of the locus is covered by the 'Unannotated' state, otherwise, the chromatin state with second highest coverage is called.

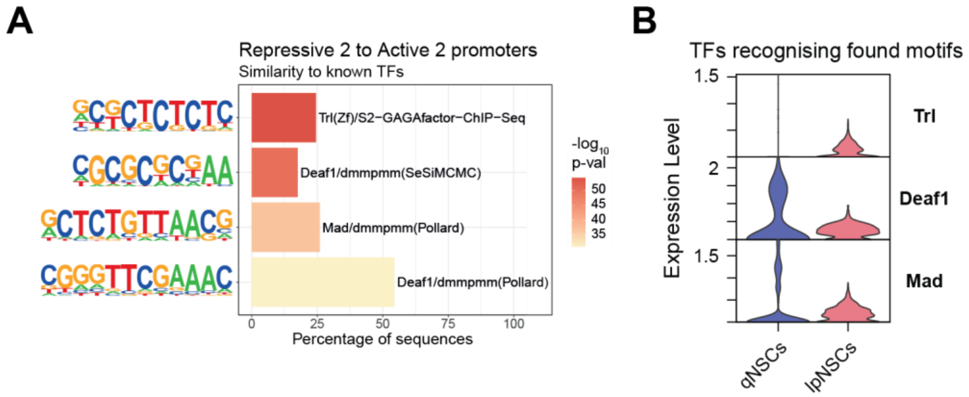


**Supplementary Figure 4. Motif enrichment analysis of promoters of genes undergoing Repressive state 2 to Active state 2 transition.** (**A**) Selected motifs identified within sequences of the promoters of the gene group in Fig. 4A and the transcription factors associated with these motifs. Promoters have been assigned as (**B**) Violin plot of the expression level of three identified transcription factors (Trl, Deaf1 and Mad) in qNSCs and lpNSCs scRNA-seq clusters. All gene expression results are based on scRNA-seq dataset from[43].
